# Cellulosic wall thickenings restrict cell expansion to shape the 3D puzzle sclereids of the walnut shell

**DOI:** 10.1101/2020.11.20.390906

**Authors:** Sebastian J. Antreich, Nannan Xiao, Jessica C. Huss, Notburga Gierlinger

## Abstract

Walnut (*Juglans regia*) kernels are protected by a tough shell consisting of polylobate sclereids that interlock into a 3D puzzle. The shape transformations from isodiametric to lobed cells is well documented for 2D pavement cells, but not for 3D puzzle sclereids. Here, we tackle the morphogenesis of these cells by using a combination of different imaging techniques. Serial face-microtomy enabled us to reconstruct tissue growth of whole walnut fruits in 3D and serial block face-scanning electron microscopy exposed cell shapes and their transformation in 3D during shell tissue development. In combination with Raman and fluorescence microscopy we revealed multiple loops of cellulosic thickenings in cell walls, acting as stiff restrictions during cell expansion and leading to the lobed cell shape. Our findings contribute to a better understanding of the 3D shape transformation of polylobate sclereids and the role of pectin and cellulose within this process.

## Introduction

Fruits of the Persian walnut (*J. regia*) are composed of a green and fleshy husk (fused bract and bracteoles), a dry and hard shell (pericarp), and a tasty and healthy kernel protected by those two envelopes. A closer look into the shell reveals polylobate sclereid cells tightly interlocked in 3D with their neighbours, which leads to a higher contact area between cells and superior mechanical properties compared to tissues with isodiametric cells like in pine seed coats (Antreich et al. 2019). Furthermore, the irregularly shaped cells are also found in shells of pecan and pistachio (Huss et al. 2020). The morphogenesis of such shell tissues is controlled by physical forces as well as biochemical signalling (Landrein & Ingram, 2019). So, using only one cell type may simplify the coordination of growth of the tissue compared to shells with a layered arrangement of different tissues found in Macadamia (Schüler et al. 2014), which make the coordination of growth more complicated.

In general, cells of plant tissues divide first and expand later during the fast growth phase of the plant organ (Gonzalez et al. 2012). During expansion, hydrostatic pressure (turgor) expands the whole cell, stretches the cell wall and forces it to loosen some parts, followed by adding new materials to grow (for a review, see Cosgrove 2018). Root and stem cells expand mainly in one axis to push the root down into the ground or the stem up into the air (Baskin 2005, Daher et al. 2018). Nevertheless, there are tissues where the cells start to expand irregularly, forming lobes like in epidermal cells of leaves (Vöfély et al. 2018). The irregular shape of the cell helps to reduce mechanical stress on the cell wall caused by high turgor pressure. For example, in growing epidermal cells of *A. thaliana*, lobes reduce the overall mechanical stresses on the cell and tissue level when cell size increases, however, high stress values become visible at the indents between the lobes (neck regions) (Sapala et al. 2018).

These irregular cell shapes generate certain stress patterns which are strongly interlinked with cell wall composition and its mechanical properties (Kierzkowski et al. 2019). The primary cell wall is composed mainly of polysaccharides like cellulose, which is the main load-bearing component; pectin, which is important for cell wall flexibility; and hemicelluloses, which cross-link cellulose microfibrils (Lampugnani et al. 2018). Cellulose is the stiffer part of the cell wall due to the microfibrillar arrangement and is linked to the cortical microtubule distribution in the cell (Bidhendi and Geitmann 2016, Gutierrez et al. 2009). These microtubules tend to orient along higher stressed cell wall regions, where more cellulose becomes deposited, thus increases the stiffness of the cell wall (Sampathkumar et al. 2014). Pectin does not only play a central role in cell-cell adhesion in the middle lamella (Marry et al. 2006), but also in lobe initialisation by changing the stiffness of the cell wall (Haas et al. 2020, Majda et al. 2017, Peaucelle et al. 2015). Recently, a two-step mechanism for lobe formation has been proposed, where de-methylated pectin increases stiffness at the future indent, which leads to cell wall undulation associated with higher stressed regions. This furthermore favours the alignment of microtubules and increased accumulation of cellulose fibrils at the indent, which slows down expansion at this location during growth (Altartouri et al. 2019, Bidhendi et al. 2019).

Most studies on irregular cell shapes focus on the epidermal pavement cells of *A. thaliana* or on epidermal cells of other dicotyledons, monocotyledons and ferns (Sotiriou et al. 2018, Vöfély et al. 2018). In the epidermis, mainly the anticlinal walls undulate, while the periclinal walls are straight, which makes it easy to measure with confocal laser scanning microscopes in 2D. Based on that, shape descriptors are also established in 2D (Poeschl et al. 2020, Sapala et al. 2018, Altartouri et al. 2019, Vöfély et al. 2018). But how do the sclereid puzzle cells form in 3D in walnut shells?

The challenge in walnut is that the husk covers the shell tissue during fruit growth and cells in the shell expand irregularly in all directions. In this study, we uncover this morphogenesis for the first time in 3D by using serial block face-scanning electron microscope (SBF-SEM). Based on the 3D reconstructions, we characterise cell shapes with different shape descriptors. We also investigate the developing sclereids with Raman spectroscopy to understand the chemical contributions to lobe formation. Finally, we suggest a possible mechanism for shaping walnut puzzle sclereids in 3D.

## Results

### Walnut and tissue growth

Our first step to track lobe formation in walnuts was a detailed monitoring of the growth and tissue development during the year of 2019. The strongest increase in weight and size occurred between 6 to 10 weeks after catkin formation (WAC), corresponding to 3^rd^ of June to 1^st^ of July, when walnut weight increased 27-times (from 1.7 ± 0.2 g to 46.5 ± 4.3 g) together with length and width (Fig. 1a). From WAC 4 to WAC 12, tissue development was reconstructed from picture stacks made by serial face-microtomy (SF-M), which reveal a strong increase of shell volume in this period (Fig. 1b-c, Supp. video. 1). In the beginning (WAC 4-6), the kernel is only presented as a small embryo, which expands fast into the already formed cavity (locule) shaped by the inner part of the shell (Supp. Fig1), until it filled this space at WAC 10. At the same time, the shell reached its final size and lignification started, initially along the suture from tip to base.

**Fig. 1.**
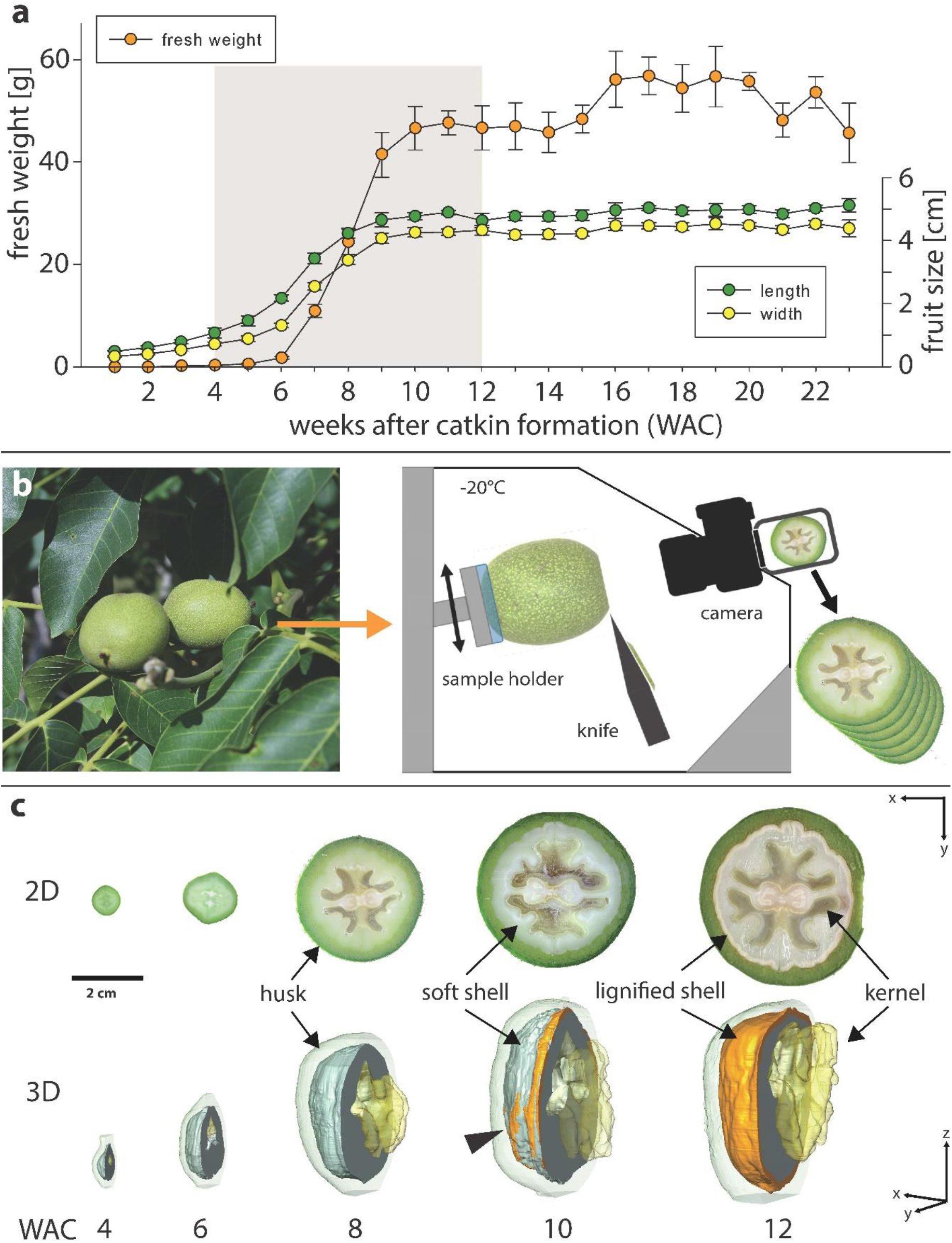
Walnut fruit development: **a)** Fresh weight, length and width of walnut samples 1-23 weeks after catkin formation (WAC), corresponding to end of April to end of September in 1-week intervals. The grey area is showing the period chosen for serial face-Microtomy (SF-M) (n = 5, error bars = SD). **b)** Freshly collected walnuts were transferred into the cryostat microtome chamber, sequentially cut and photographed. **c)** 3D reconstructions from SF-Microtomy show changes of the internal built-up of the walnut (kernel, soft shell, lignified shell and husk). Lignification starts along the suture but also appeared at some areas far away from the suture (arrowhead).

### Cell size and shape changes

During the 8 weeks of tissue growth, the cell shapes were analysed with SBF-SEM followed by 3D reconstructions (Fig. 2a). This detailed investigation showed a strong increase in cell size during the expansion phase of the (drupaceous) nut (Fig. 2b). Mainly from week 6 to week 10 cell size increased 13-fold (from 7.1 × 10^3^ μm^3^ to 94.1 × 10^3^ μm^3^). Cell surface area expanded in the same period 8-fold (from 2.2 × 10^3^ μm^2^ to 17.6 × 10^3^ μm^2^) (Supp. Fig. 2). To characterise the transition from small isodiametric cells to large polylobate cells according to reconstructions of the SBF-SEM stacks (Fig. 2c) cell shape descriptors for 3D development are introduced.

**Fig. 2.**
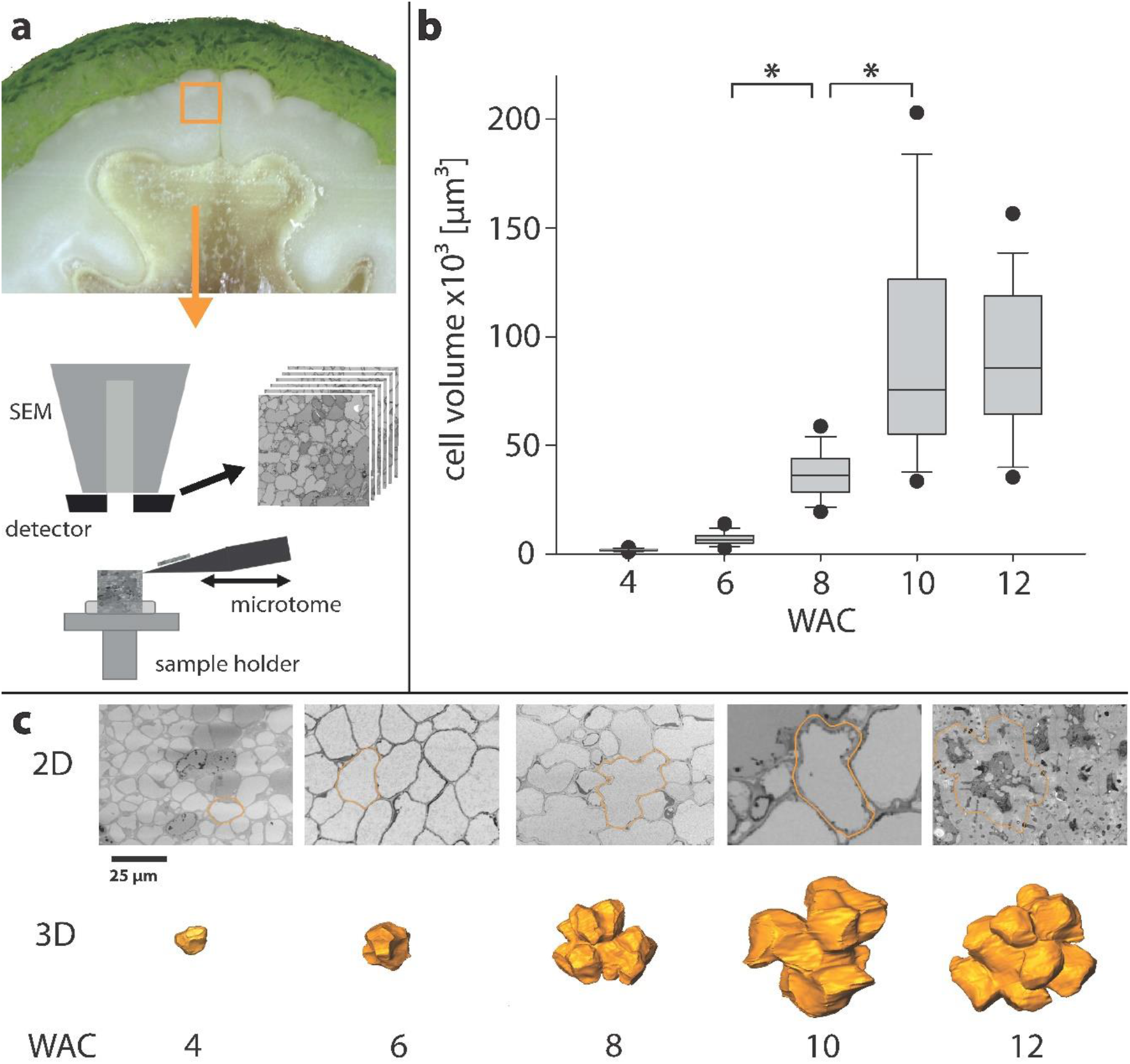
Cell size and shape development: **a)** Small pieces of shell close to the suture were cut out, fixed and embedded for the SBF-SEM to produce serial cuts. **b)** Cell volume based on reconstructions during the growing period in weeks after catkin formation (WAC) (n > 50, points = 5/95 percentile). **c)** SBF-SEM images represent each developmental stage. The cells marked on the picture have a volume closest to the average value from b) and are shown as 3D reconstruction below (scale bar is the same for all pictures).

Shape descriptors like circularity (form factor), solidity or convexity exist for 2D pavement cells of *A. thaliana* (Poeschl et al. 2020). To describe the changes of the walnut cells during development we also used solidity, which represents the ratio between cell volume and convex hull volume (Fig. 3a). The solidity was 0.84 ± 0.05 at week 4 and dropped to 0.61 ± 0.06 at week 12.

**Fig. 3.**
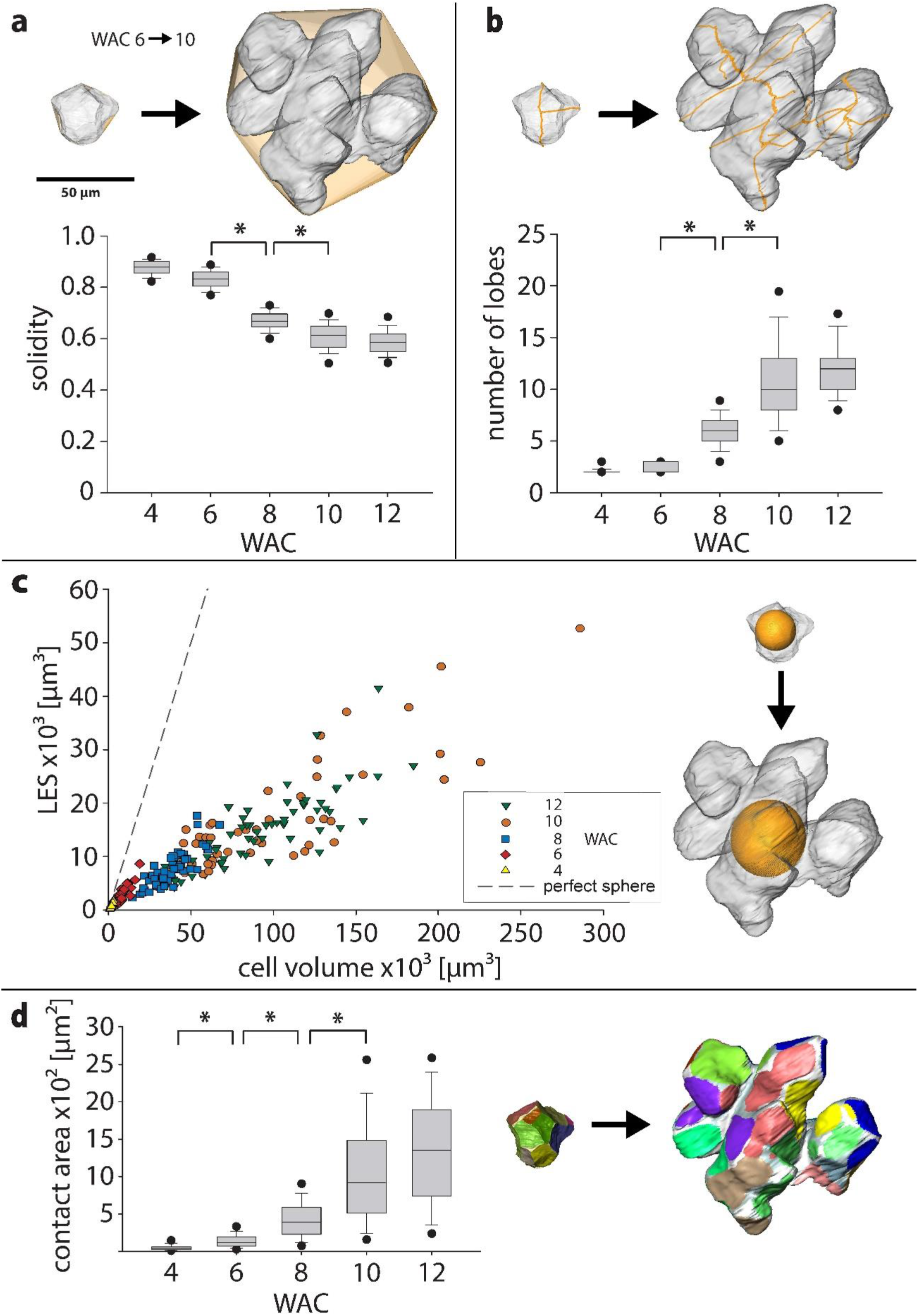
Cell shape descriptors in 3D: a-d) Changes in cell shape descriptors from week 6 to week 10 illustrated on the same set of cells (small cell from June 3^rd^, big cell from July 1^st^). During this interval we can observe **a)** a decrease of solidity, **b)** an increase in the number of lobes after skeletonization, **c)** a 5-times lower increase of the largest empty sphere (LES) compared to a hypothetical spherical cell and **d)** a strong increase of cell contact area of neighboring cells and intercellular spaces (non-colored).

Another tool to describe cell shape changes in 2D and 3D is the skeleton of the cell. The cell shape is reduced to the innermost line and the skeleton endpoints correspond to the number of lobes (Fig. 3b). During morphogenesis the main lobe number increased steadily from isodiametric cells (2 lobes) to polylobate cells with around 12 lobes.

Turgor-pressure causes the cell wall to bulge outwards, leading to mechanical stress on the cell wall (Cosgrove 2018). In pavement cells of *A. thaliana,* the largest empty circle (LEC) that fits into the cells was used as a proxy for the maximal stress on the cell wall (Sapala et al. 2018). But these cells have a relative constant vertical thickness, whereas the walnut cells expand non-uniformly in all directions during growth. To extend this factor into 3D, we introduced the largest empty sphere (LES), which describes the biggest sphere that fits into the cell volume (Fig. 3c). With growing cell volume, the LES of the growing walnut cells increased less compared to a hypothetical cell without lobes (represented by a perfect sphere). Similar to *A. thaliana*, the maximal cell wall stress during cell expansion increased around 5-times less by the formation of lobes.

With decreasing solidity, the cell became more lobed, which resulted in an increase of the cell surface area. Together with the fact that the number of cell neighbours stayed constant during development (Supp. Fig 3), cell contact area between neighbouring cells increased strongly (8-fold) from week 6 to week 10 (Fig.3d). 3D reconstruction of single cells revealed that contact areas become separated by intercellular spaces (ICS) resulting in more but smaller single areas.

### Cell wall changes

The changes in cellulose deposition were followed during the developmental period by staining microsections with calcofluor white (Fig. 4a). At WAC 8 and 10 loops of cellulose become visible, also seen by light microscopy (Supp. Video 2). Additionally, cell wall thickness of single cells in week 8 were analysed in detail in SBF-SEM reconstructions. The average thickness was 0.88 ± 0.22 μm with clearly thicker sites at the cell indents (Fig. 4b). By visualising the parts which were thicker than the average cell wall thickness (values > 0.88 μm), loops of thicker cell wall became visible (Supp. Video 3). In week 10, the average cell wall thickness doubled (1.62 ± 0.44 μm), the loops remained, but less pronounced due to the thicker walls near the indents (values > 1.62 μm) (Supp. Fig 4a, Supp. Video 4). In order to follow the chemical composition of the cell wall in context with the microstructure, Raman imaging (Gierlinger 2018) was performed on freshly cut cross-sections from week 8, focussing on the cell wall of the indents (Fig. 4c). Integration of the CH ‒stretching region from 2831-3009 cm^−1^ maps all organic materials of the region of interest. Non-negative matrix factorization (NMF) (Prats-Mateu et al. 2018) unmixed two cell wall endmembers with different chemical composition (Fig. 4d). One endmember spectrum revealed mainly cellulose signals (blue spectrum with bands at 1095 and 1380 cm^−1^ and was found in the tip of the indents (Fig. 4c). In contrast, the second endmember included clear pectin signals (Fig. 4d, green spectrum with pectin marker band at 843 cm^−1^) and was mainly found on the sides of the ICS (where the middle lamella is located) (Fig. 4c). Sections collected in week 10 confirmed also higher pectin accumulation at the corners of the ICS and a high cellulose signal at the indent and along the cell walls (Supp. Fig 4b, c).

**Fig. 4.**
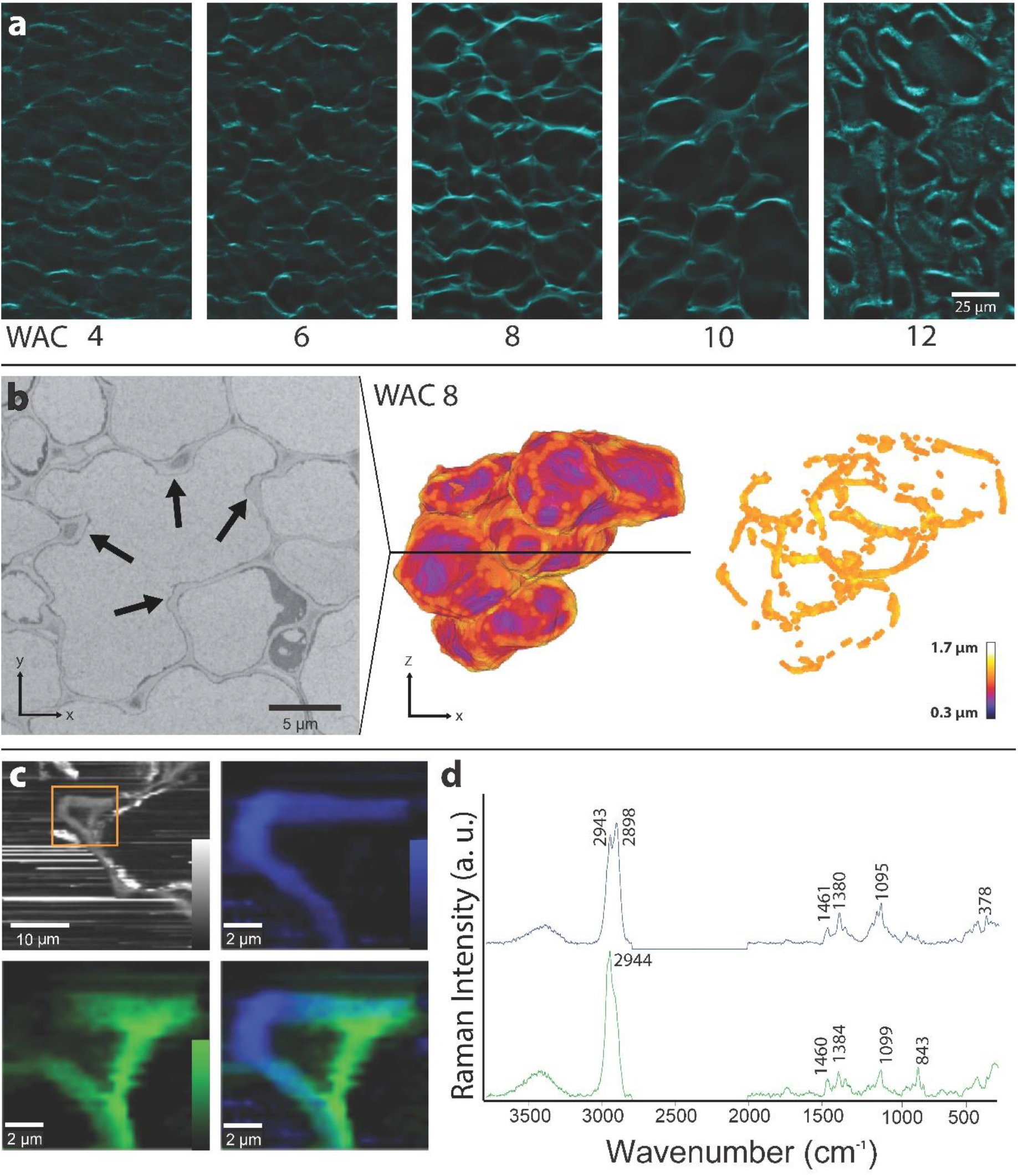
Cell wall thickening and cellulose deposition. **a)** MeOH fixed and de-colored sections of all developmental stages after calcofluor white staining show loops of cellulose all over the tissue in week 8 and week 10. In week 12, only the secondary cell wall towards the lumen is still unlignified and thus the only part which is stained. **b)** One section of the SBF-SEM stack located along the black line in the 3D model. Cell showed lobes due to several indents (arrow). The cell wall was visualized based on thickness. After removing cell wall elements, which are thinner than the average cell thickness, loops become visible. **c)** Raman imaging of a section: integrating the CH– stretching region from 2831-3009 cm^−1^ revealed the organic material of the cell wall and deposits along the cell wall. A zoom into the indent region based on non-negative matrix factorization (NMF) highlighted cellulose (blue) on the indent tip and pectin accumulation (green) on the sides and the opposite site of the ICS. **d)** The endmember spectra confirm pure cellulose on the indent (blue) and a pectin rich region (green).

## Discussion

Walnut fruits showed the strongest increase in fresh weight between end of May to mid of July (WAC 4 to 12), which is confirmed by other studies on walnut fruit development (Drossopoulos et al. 1996, Pinney and Polito 1983). Our investigation focussing on the shell development in this period revealed strong changes in cell shape - from small isodiametric to big polylobate cells. Especially between week 6 and week 10, the cells had the strongest volume and surface increase and formed the lobes.

### Lobe formation of cells of walnut shell tissue

The formation of irregular cell shapes is well studied in epidermal cells of *Arabidopsis thaliana* (Sampathkumar et al. 2014, Sapala et al. 2018, Altartouri et al. 2019, Bidhendi et al. 2019). Our findings in the shell of walnut showed similar features during development. In the beginning of development cell walls were straight between two freshly divided cells (Fig. 5a). With continuous age and size, the cell wall started to undulate which leads to a concave appearance of some contact faces to neighbouring cells (Fig. 5b). The reason for this undulation can be due to changes in stiffness of the cell wall or changes in pectin composition (Haas et al. 2020, Altartouri et al. 2019, Majda et al. 2017). At the innermost part of the concave cell wall, higher stresses caused by turgor pressure will arise, similar to the epidermal cells of *A. thaliana* (Sapala et al. 2018). However, contrary to *A. thaliana*, walnut shell cells expand non-uniformly in 3D causing loop like stress patterns. To counteract these stresses cellulose is deposited along the future indents to thicken the wall – a process that is probably mediated by cortical microtubules and leads to the observed loops of cellulose (Fig. 5c, Supp. Video 5). These cellulosic thickenings likely hinder expansion at the formed indents and the expansion of the cell toward neighbouring cell corners is promoted. The difference in expansion caused by thicker walled indents is measured by Elsner et al. (2018) in *A. thaliana*, where tip regions of indents expand slower than the side regions.

**Fig. 5.**
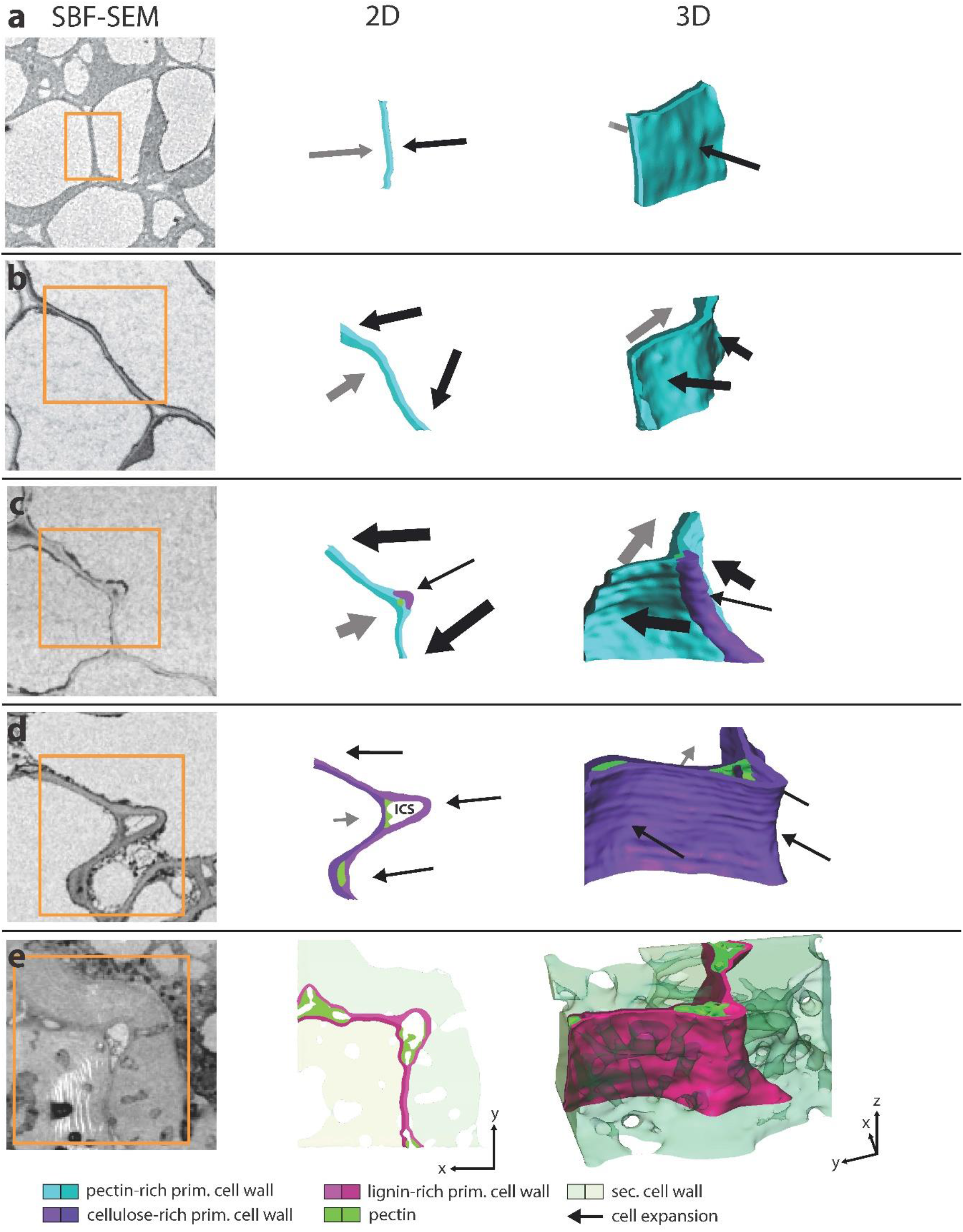
Mechanism of lobe formation in walnut shell: representative sections showing the cell wall of neighboring cells in each developmental stage (same scaling), a 2D-sketch and a 3D visualization of the same area shows the formation of the indent. **a)** After cell division the cell wall is straight and cell expansion is weak (thickness of arrows represents expansion speed and direction). **b)** The cell expands faster, and the cell wall starts to undulate, causing different expansion directions of the neighboring cells. **c)** Due to higher cell wall stresses at the curved section, cellulose gets deposited and the thickness increase at this location. The cell expands further but less strong at this position and an indent starts to form. A small intercellular space (ICS) is created, first filled completely. The opposite cell expands off-plane and forms another loop of wall thickening. **d)** Later during cell growth, the ICS gets bigger and shows an open space, sometimes exhibit protrusions. The whole cell wall becomes richer in cellulose and cells expand less strong. **e)** Finally, secondary cell wall is deposited, and the primary cell wall gets lignified, which stops the cell expansion.

In our study, we showed that the restriction of expansion was so strong that cell contacts to the neighbouring cells were lost at the indents and an ICS was formed. In the beginning the ICS was filled by strongly stained materials, later the ICS opened completely and the contact to the adjacent cell wall was lost (Fig. 5d). Raman images showed high pectin signals at the edges of the ICS close to the middle lamella. As pectin not only holds the cells together via the middle lamella but also controls the separation of cells (Daher and Braybrook 2015), especially at cell corners and along the ICS turgor mediated forces are highest (Javis 1998). At these locations high amounts of the highly de-esterified homogalacturonan are presented, which increase the viscosity of the cell wall matrix via Ca^2+^ bridges and delimit cell wall separation and ICS formation (Sotiriou et al. 2018, Giannoutsou et al. 2013, Parker et al. 2001, Knox et al. 1990).

However, in walnut, the stiff restrictions and the strong cell expansion formed new ICS all along the cells, which is more analogue to mesophyll tissue of *Zea mays* (Giannoutsou et al. 2013) or *Vigna sinensis* (Sotiriou et al. 2016) than to epidermal tissue, where cell-cell contact is continuous (Sotiriou et al. 2018). In *Z. mays,* cellulose deposition is parallel to cortical microtubule orientation, which form ring-like thickenings around the whole cell perpendicular to the leaf axis (Apostolakos et al. 1991). It is shown that during tissue expansion cells become lobed due to cellulose depositions and the resulting ICS becomes continuously bigger. The same mechanism for lobe formation can be proposed for cells of the walnut shell but, contrary to *Z. mays,* the loops of cell wall thickenings are not orientated but randomly distributed. Therefore, each individual cell shapes and gets shaped by other cells when they expand into new ICS between cells, where the walls exhibit less resistance. This leads to the observed variability of cell shapes in the shell tissue and the network-like appearance of the ICS (Supp. Fig 5). As development proceeds, cellulose was deposited along the whole cell wall, reducing the local variability in thickness and therefore the loops became less pronounced.

Cell expansion and lobe formation ended with the onset of secondary cell wall formation and incorporation of lignin into the primary cell wall, as indicated by stronger staining in SEM pictures (Fig. 5e) and is confirmed in previous studies (Antreich et al. 2019, Xiao et al. 2020).

### Lobed cell shape is beneficial for stress resistance on the cell and tissue level

The change from isodiametric to polylobate cells happens mainly within 4 weeks. All shape descriptors significantly changed within this period. More lobes are formed (more skeleton endpoints) and became more pronounced (reduced solidity), which led to a drastic increase in contact area to neighboring cells. As shown in seeds of *Portulaca oleracea,* the wavy sutural interface between neighboring cells of the seed coat increase overall strength and fracture toughness compared to straight cell interfaces (Gao et al. 2018). In the same way the interlocking of the polylobate sclereid cells in walnut lead to high values in tensile and compression tests on the tissue level (Antreich et al. 2019, Huss et al. 2020). On the cellular level, cells kept their LES low during development to reduce high stresses on the cell wall analogue to epidermal cells in *A. thaliana* (Sapala et al. 2018). So, the polylobate cell shape has two functions: on the one hand, it reduces internal stresses on the cell wall during development and, on the other hand, it increases tensile and compression strength of the whole mature shell tissue. Models derived from plant samples show that cell size and shape with its mechanical constraints influence tissue growth in 2D (Sapala et al. 2018) and 3D (Bassel et al. 2014). Additionally, the surrounding tissues could have a strong influence on tissue morphogenesis. As shown in *A. thaliana* seeds, the pressure of the endosperm and the restriction of the seed coat affect microtubule orientation and cell wall thickening of mechanosensitive cells (Creff et al. 2015, Beauzamy et al. 2016). In the case of the walnut, mechanical interactions may derive from the expanding embryo and the restricting husk forcing the cells of the shell to interlock. Under these assumptions it would be interesting to use our data to create 3D finite element models on cellular level to shed more light on the morphogenesis of the whole walnut shell tissue.

### New insights into walnut development due to 3D visualization

SF-M and SBF-SEM are promising tools to study the morphogenesis of plant organs and tissues in 3D. In our study, SF-M is a simple and cheap tool to give insights into young and soft tissues, where X-ray computer tomography methods reach their limitation regarding the loss of contrast due to water content and loss of sharpness due to movements of the sample (Kaminuma et al. 2008, Kuroki et al. 2004). Especially with samples showing differently colored tissues, the colored pictures unfold their full potential. Contrary, SBF-SEM gives insights into cell organization with impressively high resolution. Studies on microtubules of the mitotic spindle in human cells (Nixon et al. 2017) or on ER organization in *Z. mays* (Arcalis et al. 2020) are the beginning trend in 3D ultrastructure investigation (Smith & Starborg 2018). Also, in this study, SBF-SEM allowed us to analyze for the first time the shape transformation of the 3D sclereid puzzle cells in walnut shell tissue. Furthermore, complex structures like ICS network can be visualized in 3D in more detail than using casting methods (Prat et al. 1997) and is independent of gas-filled space needed for X-ray computer tomography scans (Kuroki et al. 2004). Further, SBF-SEM could be of big interest in the study of cell development in *A. thaliana* to establish life like 3D models to better understand the role of periclinal walls in the formation of undulating cell walls (Majda et al. 2017, Majda et al. 2019, Bidhendi, 2019).

Finally, the combined use of state-of-the art 3D-characterisation and micro spectroscopic methods will shed new light on still open questions, e.g., the stiffness differences in the beginning of cell wall undulation or the distribution of microtubules during lobe formation. Revealing the whole formation process of the 3D sclereid puzzle cells in walnut and comparing it with shells of other nuts will help us to understand the general concept of shell morphogenesis in plants.

## Materials and Methods

### Sampling

We collected walnuts in 1-week intervals throughout the year 2019, starting from end of April until end of September, from the horticulture garden of BOKU, Vienna. Walnuts grew on a >40-year-old tree of the cultivar ‘Geisenheim 120’. Always 5 nuts were collected from the sunny side of the tree put into plastic bags and immediately brought to the labs for further investigation.

### Fresh weight, size, serial face-microtomy (SF-M)

Each week the fresh weight, length and diameter of each nut was measured. Every two weeks (from week 4 to 12) one of the five walnuts was used for the SF-M. Another walnut was used for the Serial Block Face-Scanning Electron Microscope (SBF-SEM), calcofluor white staining and Raman microscopy analysis. All other nuts were frozen at −20°C for later use. For the SF-M the walnut was kept in the cryostat microtome (CM3050 S, Leica Biosystems, Nussloch, Germany) for 1-4h (depending on the nut size) at −20°C until all liquids in the walnut were frozen. A camera was mounted in front of the walnut and after each 30-100μm cut (depending on the walnut size) with the microtome knife a photo was made. As the sample holder moves toward the knife, the camera position needed no changing during the cutting. The acquired picture stacks of the whole nuts were processed and registered in ImageJ (NIH, Bethesda, Maryland) with the plugin ‘Linear stack alignment with SWIFT’ using the standard settings (Rueden et al. 2017). Then the aligned stack was segmented in the Software Amira (Thermo Fisher Scientific, Waltham, Massachusetts) into seed, soft shell, hard shell and husk, followed by 3D-reconstruction.

### SBF-SEM

Around 1 mm x 1 mm x 1mm small pieces of walnut shell were trimmed with a razor blade, always from the mid region of the nut close to the suture. Trimmed pieces were immersed immediately in fixation solution containing 3% glutaraldehyde in 100 mM sodium cacodylate (pH7.4) and stored at 4°C overnight. Samples were rinsed 3 times with 150 mM cacodylate buffer and post fixed with 2% osmium tetroxide and 0.2% ruthenium red in 150 mM cacodylate buffer for 1 hour at room temperature. After 5 times washing with cacodylate buffer, samples were incubated in freshly prepared thiocarbohydrazide solution (1% w/v in dH2O) for 45 minutes, followed by 3 times washing with dH2O and post-fixed a second time with a 2% osmium solution for 1 hour. Samples were washed again 4 times with dH2O immersed in 0.5% uranyl acetate and stored overnight at 4°C. Again, samples were washed 5 times in dH2O and then transferred in Waltron’s lead aspartate solution for 30 min at 65°C, followed by 5 times washing in dH2O. Dehydration was performed in 30%, 50%, 70%, 90%, 100%, 100% ethanol in water, followed by 100%, 100% acetone; each 30 minutes at room temperature. Samples were then infiltrated by 25% low-viscosity resin in acetone and left at 4°C overnight. Then samples were transferred into 50% and further into 75% resin, 4 hours each, until 100% resin overnight at 4°C, followed by a second round of 100% resin for 6h at room temperature. Samples were then embedded in flat embedding moulds and polymerised at 65°C for 48 hours. Resin blocks were trimmed with a glass knife on a UC-7 ultramicrotome (Leica Microsystems, Vienna, Austria) to 0.5 mm^3^ and glued with silver cement on a stub. They were coated with a 10 nm gold layer in an EM SCD005 sputter coater (Leica Microsystems, Vienna, Austria) and mounted on the microtome of the VolumeScope SEM (Thermo Fisher Scientific, Waltham, Massachusetts). Scans of 100 μm^2^ were acquired with 1.18 kV, 100 pA, and 3 μs dwell time. Approximately 1000 slices with a slicing depth of 100 nm were made, controlled by the software Maps 3.4 VS (Thermo Fisher Scientific, Waltham, Massachusetts). The resulting stacks were scaled to a useable size (around 1000 × 1000 × 100 px) for the Amira software and registered in ImageJ with the plugin ‘Linear stack alignment with SWIFT’ using the standard settings. All whole cells were segmented manually, which were not cut off by the border. From each segmented cell surface/volume, convex hull surface/volume and contact surfaces between each neighbouring cell was calculated in the software Amira. Additionally, lobe number was calculated by the centreline tree function (tube parameter: slope: 1.2, zeroVal 3.5). This made a skeleton of the cells in 3D but was very sensitive to rough cell shape. Therefore, the segmented cells were smoothed to eliminate selection artefacts, so that only main lobes were counted. Finally, the largest empty sphere (LES) of each cell was calculated with the ‘Thickness’ function of the ImageJ plugin BoneJ (Dougherty & Kunzelmann, 2007). 3D-reconstraction of all cells (inclusively cell shape descriptors) were done in the software Amira.

### Confocal Laser Scanning Microscopy (CLSM)

Small pieces of shell tissue were cut out close to suture and fixed and de-coloured according to Pasternak et al. (2015) with minor changes to stain cellulose with calcofluor white. Samples were put into an Eppendorf tube with 1.5 ml pure MeOH for 20 min at 37°C. Afterwards the sample was transferred into 0.8 ml fresh pure MeOH for another 3 min, then 200 μl dH2O was added in 2 min intervals until reaching 2 ml in total. After this, samples were washed twice with dH2O for 5 min each. Afterwards samples were transferred on a glass slide, stained with one drop of a ready-to use calcofluor white stain solution (Sigma-Aldrich) containing 1 g/l calcofluor white M2R and 0.5 g/l evans blue and then mounted on a TCS SP5 II CLSM (Leica Microsystems, Vienna, Austria). As emission source a 405 nm UV diode was used, and detection range was set from 450 to 500 nm. Pictures were made with the same magnification using a 40x/0.85 objective and a resolution of 0.2 μm.

### Confocal Raman Microscopy

From small blocks of frozen walnut shells 20-30 μm thin sections were cut in the cryostat microtome and transferred on a standard glass slide. Samples were washed several times with dH2O, followed by D2O and sealed with nail polish for Raman microscopic measurements. Spectra were acquired from micro sections using a confocal Raman microscope (alpha300RA, WITec, Ulm, Germany) equipped with a 100× oil immersion objective (NA 1.4, Carl Zeiss, Jena, Germany) and a piezoelectric scan stage. A laser (λ = 532 nm) was passed through a polarization-preserving single-mode optical fibre and focused through the objective with a spatial resolution of 0.3 μm on the sample. The Raman scattering signal was detected by a CCD camera (Andor DV401 BV, Belfast) behind a spectrometer (600 g mm^−1^ grating, UHTS 300 WITec, Ulm, Germany). The laser power was 40mW. For measurement setup the software Control Four (WITec, Ulm, Germany) was used. Raman analysis was performed with Project FOUR (WITec, Ulm, Germany) and Opus 7.5 software (Bruker Optik GmbH, Ettlingen, Germany). After applying cosmic ray spike removal, Raman chemical images were generated based on the integration of relevant wavenumber regions (e.g., CH stretching). The indent was selected, and a non-negative matrix factorization (NMF) was performed in Project FOUR with six basis spectra.

### Statistics

Data were analysed with the software SigmaPlot 12 (Systat Software, San Jose, California) for significant differences between each development stage. On all data from the cell segmentation a Kruskal-Wallis one-way analysis of variances on ranks was performed followed by Dunn’s Method to compare all ranks. Significant differences (p<0.05) were marked in the figures with *.

## Acknowledgements

The authors thank the whole BIONAMI group for all the help and comments. We also thank Karl Refenner for giving us access to the walnut tree on the horticultural garden of BOKU, Vienna. Thanks to Ingeborg Lang for critical reading of the manuscript and to Elsa Arcalis and Ulrike Hörmann-Dietrich for providing chemicals and lab space for the SBF-SEM sample preparation. The authors acknowledge funding from the European Research Council (ERC) under the European Union’s Horizon 2020 research and innovation program grant agreement No 681885 and the HSRM Project NANOBILD for infrastructure support.

