## Supplemental data for "Cellulosic wall thickenings restrict cell expansion to shape the 3D puzzle sclereids of the walnut shell"

**Supporting data**

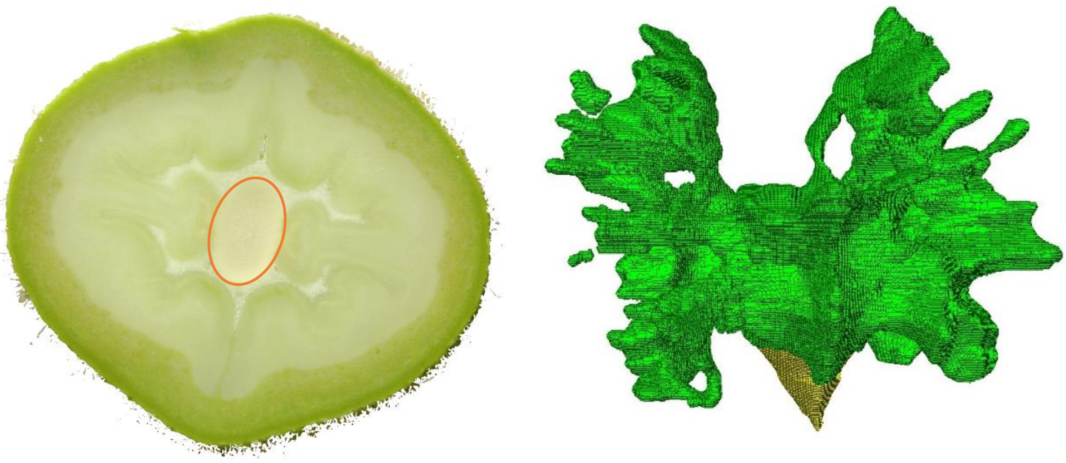

**Supp. Fig 1 walnut cavity: a)** in week 6 the embryo of the walnut (marked orange) exhibit only a small space of the whole cavity formed by the surrounding shell tissue. **b)** 3D representation of the embryo (yellow) and the formed cavity (green)

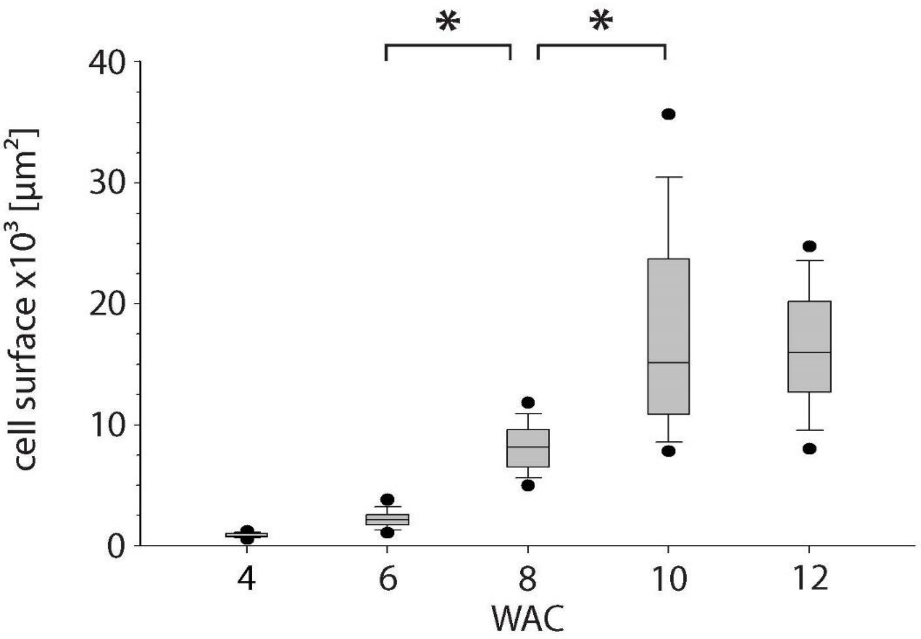


**Supp. Fig 2 cell surface**: cell surface of at least 50 cells from week 4 to week 12 after catkin formation (WAC). Here the same trend as in the cell volume is visible (n > 50, points = 5/95 percentile).

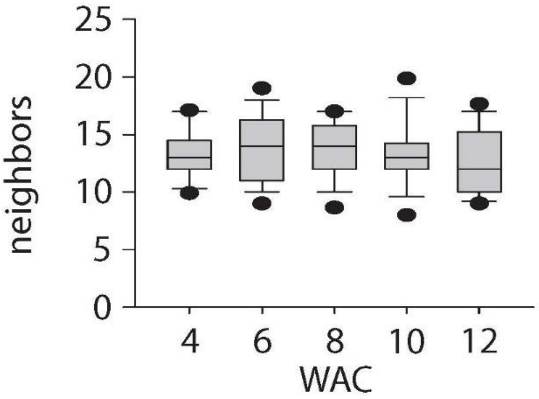


**Supp. Fig 3 cell neighbors:** average number of cell neighbors counted after 3D segmentation

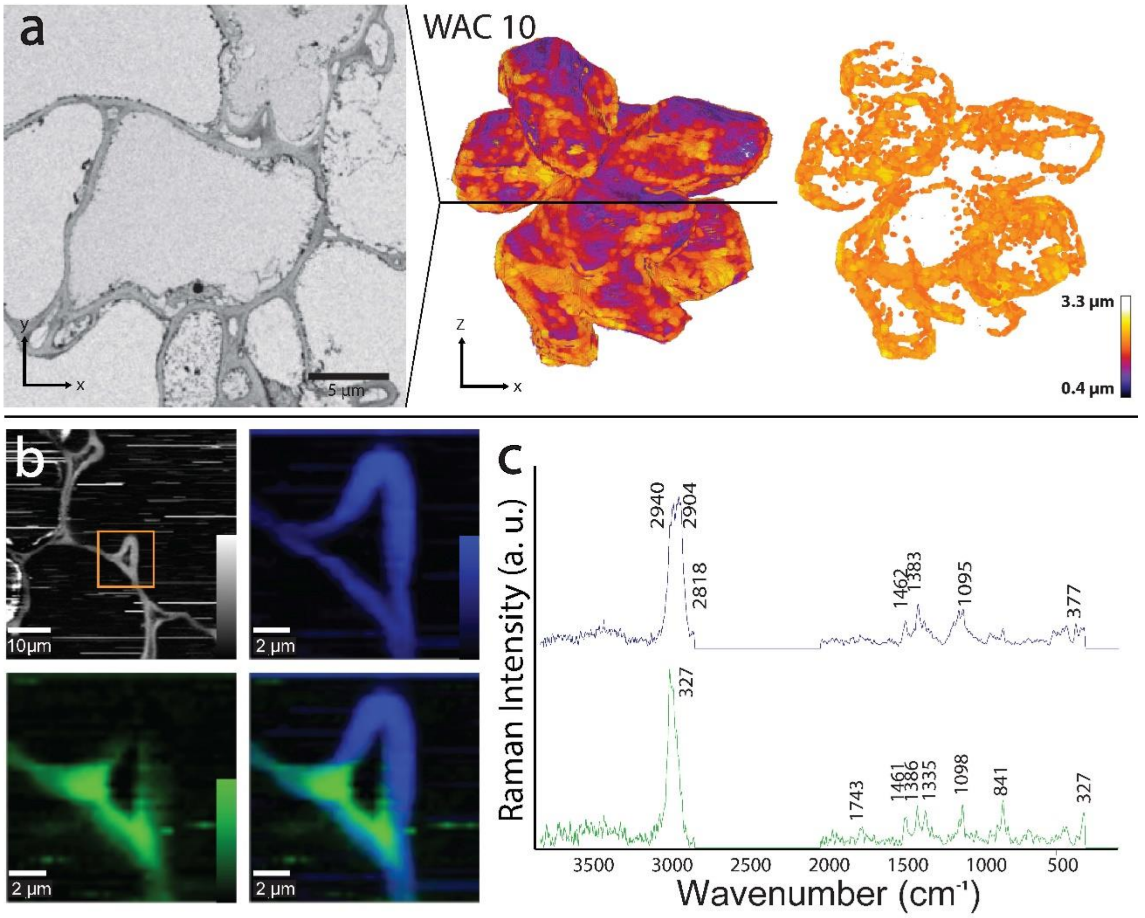


**Supp. Fig 4 Raman imaging analysis on the indents at week 10: a)** one section of the SBF-SEM stack located along the black line in the 3D model. The cell wall was selected and the thickness was visualized. After removing cell wall elements, which are thinner than the average cell thickness less pronounced loops are visible. **b)** Raman imaging of a section integrating the CH–stretching region from 2831-3009 $cm^{-1}$ reveals the organic material of the cell wall is uniformly distributed, surrounded by lipids. A zoom into indent based on non-negative matrix factorization (NMF) highlight cellulose (blue) on the indent and along the cell wall (also of opposite cell). Pectin accumulation (green) on the sides of the ICS. **c)** the endmember spectra confirm pure cellulose on the indent and the cell wall (blue) and a pectin rich region (green).

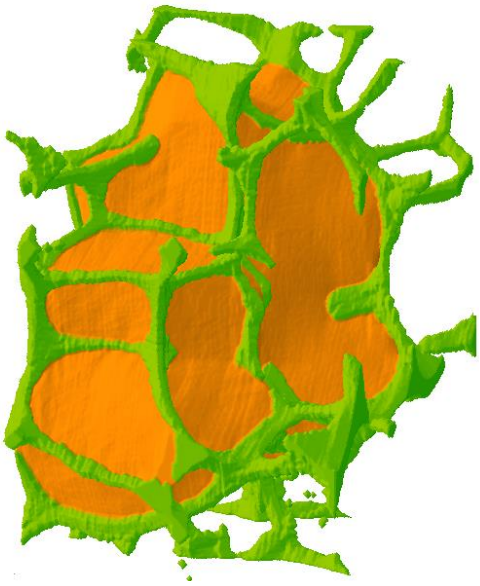

**Supp. Fig 5 intercellular space (ICS):** 3D reconstruction of the ICS (green) around a single cell (orange) in week 12 showing a net-like shape which is continuously distributed between the cells of the shell tissue

**Supp. Video 1 Serial face-microtomy:** Serial cut through the whole walnut fruit of week 6 and 3D reconstruction of the different tissues (husk, soft shell and kernel) after segmentation.

**Supp. Video 2 Loops formation in week 8:** Light microscope stack through a tissue from week 8 after de-coloration but before calcofluor white staining showing the looped cell wall thickenings.

**Supp. Video 3 Cell thickenings along the indents:** 3D reconstruction of a single cell from week 8. The cell wall was selected and the thickness was visualized. After removing cell wall elements, which are thinner than the average cell thickness loops are visible.

**Supp. Video 4 Cell thickenings along the indents and the cell walls:** 3D reconstruction of a single cell from week 10. The cell wall was selected and the thickness was visualized. After removing cell wall elements, which are thinner than the average cell thickness loops are visible but less pronounced than in week 8.

**Supp. Video 5 Serial block face-scanning electron microscopy:** Serial cut of a cell wall with an indent between two neighboring cells and 3D reconstruction of the formed cell wall thickening along the indent.
